# TriPepSVM - *de novo* prediction of RNA-binding proteins based on short amino acid motifs

**DOI:** 10.1101/466151

**Authors:** Annkatrin Bressin, Roman Schulte-Sasse, Davide Figini, Erika C Urdaneta, Benedikt M Beckmann, Annalisa Marsico

## Abstract

In recent years hundreds of novel RNA-binding proteins (RBPs) have been identified leading to the discovery of novel RNA-binding domains (RBDs). Furthermore, unstructured or disordered low-complexity regions of RBPs have been identified to play an important role in interactions with nucleic acids. However, these advances in understanding RBPs are limited mainly to eukaryotic species and we only have limited tools to faithfully predict RNA-binders from bacteria. Here, we describe a support vector machine (SVM)-based method, called TriPepSVM, for the classification of RNA-binding proteins and non-RBPs. TriPepSVM applies string kernels to directly handle protein sequences using tri-peptide frequencies. Testing the method in human and bacteria, we find that several RBP-enriched tripeptides occur more often in structurally disordered regions of RBPs. TriPepSVM outperforms existing applications, which consider classical structural features of RNA-binding or homology, in the task of RBP prediction in both human and bacteria. Finally, we predict 66 novel RBPs in *Salmonella* Typhimurium and validate the bacterial proteins ClpX, DnaJ and UbiG to associate with RNA in vivo.

## Introduction

Gene regulation in eukaryotes occurs at several levels and involves the action of transcription factors, chromatin, RNA-binding proteins (RBPs) and other RNAs. RBPs and messenger RNAs (mRNAs) form ribonucleoprotein complexes (RNPs) by dynamic, transient interactions, which control different steps in RNA metabolism, such as RNA stability, degradation, splicing and polyadenylation. Numerous diseases, such as neuropathies, cancer and metabolic disorders, have also been linked to defects in RBPs expression and function (1–3).

Technological advances such as RNA Interactome Capture (RIC) has enabled proteome-wide identifications of RBPs (4). RIC utilizes UV cross-linking to induce stable RNA-protein interactions in living cells, followed by poly(A) RNA selection via magnetic oligo d(T) beads and subsequent protein identification by mass-spectrometry. RIC studies yielded hundreds of novel RBPs in e.g. human HeLa (5), HEK293 (6), Huh-7 (7) and in K562 (8) cells but also in worm and yeast (7, 9), which do not harbor canonical RBDs, as well as factors which were not previously associated with RNA biology. Among them we find enzymes, cell cycle regulators and dual specificity DNA-RNA binders, including transcription factor and chromatin components (3). The discovery of these unconventional RBPs without known RNA-binding motifs suggests the existence of new modes of RNA binding and the involvement of RBPs in previously unexplored biological processes (10).

Intrinsically disordered regions (IDRs) are widespread in the proteome and have been shown to be involved in regulatory functions, including direct RNA binding (11). RBPs identified by RIC are highly enriched in disordered regions compared to the whole human proteome and are characterized by low complexity, repetitive amino acid sequences. In particular, a low content of bulky hydrophobic amino acids and a prevalence of small, polar and charged amino acids are found in unstructured regions of RBPs. These amino acids, such as glycine (G), arginine (R) and lysine (K), as well as the aromatic residue tyrosine (Y), form shared sequence patterns among RBPs. For example RGG box binding motifs and glycine/tyrosine boxes YGG are broadly used platforms for RNA binding and can work alone or in combination with classical RBDs (11).

The occurrence of IDRs within RBPs appears to be conserved from yeast to humans. In earlier work, we defined a core set of RBPs conserved from yeast to human and identified [K]- and [R]-rich tripeptide repeat motifs as conserved across evolution, making IDRs plastic components of RBP co-evolution (7). Importantly, we proposed that the number of repeats in IDRs of RBPs considerably expands from yeast to human, while the number of RBDs remains the same, linking repeat motifs in IDRs of RBPs to the functional complexity of regulation in higher eukaryotes.

Research over the past two decades has revealed extensive post-transcriptional control in bacteria as well, with regulatory networks comprising RBPs and small non-coding RNAs (sRNAs). Advances in RNA sequencing methods have enabled the discovery of new bacterial RBPs and showed that bacterial RBPs are relatively simple, often possessing only a few or even a single RBD per protein, which recognizes a short RNA sequence (12). This is unlike eukaryotic RBPs, whose modular architecture enables versatility and combinatorial RBP-RNA interactions. Classical RBDs, such as the S1 domain, cold-shock domain, Sm and Sm-like domains, the double-stranded RNA binding domain, the K-homology domain and others, are widespread among bacteria (12). One third of annotated bacterial RBPs are ribosomal proteins. Other bacterial RBPs are involved in regulatory functions of transcription termination, RNA synthesis, modification and translation, like the well-known Hfq protein. The latter associates to sRNAs and leads to translational inhibition of the targeted mRNA (13).

Since interactome capture relies on the use of oligo(dT) to isolate proteins bound to mRNA, the method is not applicable to bacterial species which generally lack polyadenylation (except for degradation purposes). Although experimental approaches have been developed recently to extend RBP catalogs and to include RBPs not identified by RIC techniques (14, 15), bacterial RBPs remain poorly annotated. Therefore, computational approaches able to predict new RBPs in both pro- and eukaryotes in the absence of experimental data and in a proteome-wide manner are in high demand, in order to narrow down the list of putative candidate RBPs for further experimental investigation. Several *in silico* methods have been developed to predict RBPs from primary sequence and/or protein structure (16–19). Approaches that characterize RBPs by predicting RNA binding residues from known protein-RNA structures are computationally expensive and can be trained only on a small subset of RBPs with known structure. Such methods do not generalize well to proteins whose structure is still unknown, which is the case for most bacterial RBPs.

Computational prediction tools such as SPOT-Seq (16), RNApred (20), RBPPred (18), catRAPID (17) and APRICOT (19), either derive sequence-structure features such as biochemical properties and evolutionary conservation and train a supervised classifier to distinguish RBPs from non-RBPs or classify proteins based on whether they harbor a known RBD. Gerstberger et al. define a set of 1542 human RBPs based on whether those proteins harbor known RBDs or other RNA binding motifs via both computational analysis and manual curation (1). However, *in silico* methods that identify RBPs based on known domains might have some limitations, as one third of RBPs from recent experimental studies do not have prior RNA-binding related homology or domain annotation. Therefore such methods might generate a high percentage of false negatives, i.e. RNA-binders which lack an RBD and therefore are not predicted as such, as well as false positives, RBPs with classified RBDs that perform non-RNA binding functions (3). A recently developed method, SONAR, adopts a completely different approach for RBP prediction and exploits the fact that proteins that interact with many others RBPs from a protein-protein interaction (PPI) network are more likely to be RBPs themselves (21). Although SONAR is also suitable for the prediction of ‘unconventional’ RBPs, it heavily depends on the quality and depth of the underlying PPI network and might generate false positive predictions for those proteins that interact with other RBPs but are not RNA-binding themselves.

The many newly characterized RBPs, their nontypical RBDs (or the lack of them), as well as the observation that IDRs are subject to strong sequence constraints under the form of conserved amino acid triplets conserved in the RBPs, prompted us to explore the possibility that RBPs might be confidently predicted based purely on the occurrence of short amino acid k-mers.

We set up to predict whether a protein is likely to be an RBP or not based on primary sequence using a string kernel (spectrum kernel) support vector machine (SVM). The spectrum kernel in combination with SVMs was first successfully introduced by Leslie et al. in the context of protein family classification (22). Support vector machine classifiers have been used successfully in other biological tasks, for example the identification of specific regulatory sequences (23) and in RBP prediction as well, for example in RBPPred, which creates a feature representation based on several properties of the protein.

In this paper we describe our newly developed RBP prediction method TriPepSVM. It applies for the first time string kernel support vector machines for RBP prediction. The model uses exclusively k-mer counts to classify RBPs versus non-RBPs in potentially any species. It does not depend on any prior biological information such as RBD annotation or structure, and therefore allows RBP prediction in an unbiased manner. We show that TriPepSVM performs better than other methods in RBP prediction in human and the bacterium *Salmonella* Typhimurium. Our method recovers RBPs characterized previously in RIC studies with high sensitivity and finds both RBPs adopting classical RNA-binding architectures as well as RBPs lacking known RBDs.

## Materials and Methods

### Data sets

In the following we describe the data collection pipeline to derive annotated RNA-binding proteins and non-RNA-binding proteins, which will constitute our positive and negative data sets to train and evaluate TriPepSVM on human and *Salmonella*. Our developed collection pipeline is based on the UniprotKB database (24) and derives data automatically for a given taxon (see Supplementary Figure S1). We used the taxon identifier 9606 to collect data for *Homo sapiens* and taxon identifier 590 for the *Salmonella* clade.

#### Collection of annotated RNA-binding proteins

We utilize the gene ontology database QuickGO (25) to collect annotated RBPs from UniprotKB. We apply the term *RNA*-*binding* (GO:0003723), also including all associated sub-terms, i.e. *tRNA binding*, *snRNA binding* and *poly(A) RNA binding.* The number of annotated RBPs in the QuickGO database is limited for some organisms, therefore our pipeline also supports a *recursive mode* to collect positive data from all members of a specified taxon or branch. For example, taxon 590 will retrieve annotation for all *Salmonella* strains. In order to avoid introducing bias in our model from duplicate annotations or paralogs in UniprotKB, the software CD-Hit (26) was used to remove proteins with a sequence similarity higher than 90%.

#### Collection of non-RNA-bindig proteins

Since it is challenging to define a non-RNA-binding protein, we developed a strict filtering to generate the negative set for our method. We first collect the whole Swiss-Prot proteome of a given taxon and then remove all nucleotide-binding proteins in a step-wise manner. First, we remove ambiguous proteins with an amino acid sequence length smaller than 50 AA or greater than 6000 AA. Secondly, we utilize the Uniprot keyword database and QuickGO annotations to remove annotated nucleotide-binding proteins (adopted from (27); see full list in Supplementary Table S4 and Table S5). Finally, we discard proteins containing at least one annotated or potential RBD from collected Pfam (28) domains (see Supplementary Table S6). Similarly to the positive set, CD-Hit was used to remove redundant protein sequences at 90% sequence identity.

#### Independent validation set from RIC studies

We collected a set of experimentally confirmed RBPs from three independent interactome capture studies (6–8) and from a review on RBPs (3) to evaluate the sensitivity of our model in human. First, we excluded protein sequences that were already present in our training data set. Secondly, we evaluated TriPepSVM on the human proteome independently for all four data sets and then on their union. The sensitivity was computed as the fraction of experimentally detected RBPs which are also predicted by our model.

### TriPepSVM prediction model

TriPepSVM is a discriminative machine learning model based on Support Vector Machines (SVMs) which is trained to classify RBPs versus non-RBPs based on sequence content alone. SVMs are a class of supervised learning algorithms which, given a set of labelled training vectors (positive and negative input examples) learn a linear decision boundary by finding the optimal hyperplane to discriminate between the two classes (29). The result is a linear classification rule that can be used to classify new test points, in our case new protein sequences, into one of the two classes, RBP or non-RBP. When using a kernel in conjunction with an SVM, input points are implicitly mapped into a high-dimensional vector space where coordinates are given by feature values (22). The SVM produces then a linear decision boundary in this high-dimensional space. In our model we use the spectrum kernel for classification, a linear kernel that allows the application of SVMs to strings (and therefore to amino acid sequences). Given a number *k* >= 1, the *k*-spectrum of an input sequence is the set of all the *k*-length (continuous) sequences, also called *k*-mers, that it contains. The high-dimensional feature representation for a sequence *x*, Φ(*x*), is then a vector where each entry counts the number of times each *k*-mer, from a pool of |Σ|^*k*^ possible k-mers from an alphabet of size Σ, occurs in the given sequence. The *k*-spectrum kernel is then the dot product between two sequence feature vectors:

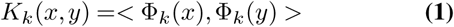

#### Model training

We randomly split the collected data into training (90%) and testing (10%) data sets, and stratified the data such that training and test data contain both roughly the same ratio of positives to negatives. As both, training and test sets are heavily imbalanced, i.e. they contain many more negatives than positives, we chose a class-weighted SVM (30) approach, which accounts for class imbalance by weighting the cost parameter *C* differently for positive and negative data points. The class-weighted SVM was used together with the spectrum kernel from the *KeBABS*-package in *R* (31). To obtain the optimal values of the hyperparamters *k*, the sub-string length and *C*, the SVM cost parameter, we perform Cross Validation (CV, *tuning* function from the *KeBABS*-package). CV splits the data into *n* equally sized subsets. This is followed by *n* iterations in which *n* – 1 splits are used for training and the *i* split is used as validation set to compute the classifier performance (see Supplementary Figure S2). Since the training of the class weights, denoted with *W* + and *W* –, is not supported in the *KeBABS*-package, we run a separate “outer” loop to select the optimal combination of weights for the positive and the negative class. In other words, for each tested combination of *W* + and *W* – we test a range of values for *k* and *C* in the CV loop and select at the end the combination of hyper-parameters which maximizes the model performance; here the model balanced accuracy (see Supplementary Paragraph 1.3.3).

#### Feature importance scores

Since the spectrum kernel *ϕ* is a linear kernel, we can obtain meaningful feature weights from the solution of the SVM optimization problem, similar to a linear regression scenario. The weight vector ***w*** for all *k*-*mers*, whose entries are the single *k-mer* contributions to the classification problem can be computed as:

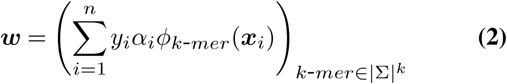

for *n* points (***x_i_***, *y_i_*), where *x_i_* ∈ ℝ^*m*^ is a m-dimensional data point and *y_i_* ∈ {0,1} is the associated label. The values of ***α*** are the results from the SVM optimization problem, and contain non-zero values for important data points, the so-called *support vectors*, which are the points closest to the SVM decision boundary (22).

Important *k*-*mers* show large absolute values in ***w*** and the sign of the weight indicates to which class, positive or negative, a *k*-*mer* contributes to.

### Application of existing methods for RBP classification

We apply different RBP prediction tools for the performance comparisons. We focus on approaches that allow for proteome-wide predictions, such as SPOT-Seq-RNA, RNAPred and RBPPred. We excluded catRAPID because the web-server does not allow submission of more than 100 protein sequences, making proteome-wide predictions unfeasible in this setting.

#### Pfam-Domain-Recognition

We collect profile Hidden Markov Models (HMMs) for known RBDs from the Pfam database (PFAM 27.0). We obtain 219 different HMMs (see bold entries in Supplementary Table S6) annotated as ‘RNA binding/recognition’ in the Pfam description, quickGO or Protein Data Bank (32). We then use the HMMER software (33) for scanning the human/*Salmonella* proteom for proteins containing at least one of the collected RBDs. We run HMMER with default parameters, i.e. using an E-value cutoff ⩽ 10.

#### SPOT-Seq-RNA

SPOT-Seq-RNA is a template-based technique to predict RNA-binding potential of a query protein (16). It employs a library of non-redundant protein-RNA complex structures and attempts to match a query sequence to the protein structure in the protein-RNA complexes by fold recognition. More specifically, SPOT-Seq-RNA requires a protein sequence in FASTA format, it passes it to PSI-BLAST to search for homologous sequences and to generate a position-specific substitution matrix (PSSM). The PSSM is used to predict several structural properties and the structural profile is matched against the known templates to compute a matching score (Z-score). Statistically significant matching templates with low free energy of the RBP-RNA complex are then used in order to assign a putative RNA binding function to the query protein. We used a local version of the tool, with *E*-*value* < 0.001 from PSI-BLAST, a minimum template matching Z-score of at least 8.04 and maximum binding free energy of −0.565, as proposed by the authors.

#### RNApred

RNApred is an SVM-based approach to predict RBPs (20) and its application supports three different modes based on: (i) amino acid composition only, (ii) evolutionary information under the form of a PSSM built from PSI-BLAST and (iii) a combination of the previous two modes with some additional refinements. Unfortunately mode (i) is the only one that can be efficiently applied proteome-wide from the RNApred web server (as the other two modes only supported submission of one protein sequence per time) and therefore this mode was chosen to apply to our test data set.

#### RBPPred

RBPPred is also based on Support Vector Machines for the classification task of RBPs versus non-RBPs (18). Compared to the other earlier tools it uses comprehensive feature representation which includes protein physiochemical properties, as well as evolutionary information under the form of a PSSM derived from PSI-BLAST, similarly to SPOT-Seq-RNA. We downloaded the command line version of RBPPred from the authors’ website and applied it on our test data set.

### Performance metrics

The performance of all methods was evaluated using two metrics: the area under the receiving operating characteristics curve (AUROC) and the area under the precision-recall curve (AUPR). Both metrics require a set of proteins which have been scored according to their likelihood of being an RBP or not, as well as the known protein class. Most of the evaluated methods, except for SPOT-Seq-RNA, return a score or probability for each protein to be RNA-binding and therefore could be assessed via these two metrics.

ROC curves plot the true positive rate versus the false positive rate of the classifier across all possible score values used to decide whether a protein is an RBP or not. The area under this curve is used to assess the classifier performance: values near 0.5 indicate that the classifier performs randomly, and an AUROC of 1.0 corresponds to a perfect classifier. However, ROC curves are known to be insensitive to class imbalance (34), i.e. the ratio of RBPs versus non-RBPs in the genome, making it hard to control the number of false positives. PR curves, which plot the fraction of predicted RBPs that are true (precision) versus the fraction of all true RBPs among all predicted RBPs (recall) at different score cutoffs, are especially suited in the latter scenario because they account for class imbalance. This gives us a better idea of the false discovery rate (1 – *precision*) of the method at all thresholds and whether the proteome-wide predictions of each classifier are likely to contain many false positives.

We also computed sensitivity, specificity, precision, balanced accuracy and Matthews correlation coefficient (35) (MCC, see Supplementary Figure S4) for the different methods, using the optimal cutoff to distinguish RBPs from non-RBPs from the PR curve, as described in Supplementary Paragraph 1.4.

### Classification of tripeptides in ordered/disordered protein regions

We analysed the relative abundance of all tripeptides in structurally disordered versus ordered regions of RBPs with the IUPred tool (36), which allows to characterize disordered protein regions lacking well-defined tertiary structure. IUPred provides a mode to predict globular domains with an average disorder score smaller than 0.5. We used this mode to classify each amino acid as ‘structured’ if it occurred in a predicted domain and disordered if it did not. From there, we were able to estimate the structural disorder fraction of each tripeptide defined as the number of times that the tripeptide was found in disordered regions of RBPs divided by the number of times the same tripeptide was found in RBPs.

### Cell culture and strains

*Salmonella enterica subsp. enterica* serovar Typhimurium strain SL1344 was cultivated in standard LB medium if not stated otherwise. We generated chromosomal insertions of FLAG-encoding sequences downstream of candidate genes using the Lambda Red technique (37, 38). For details see Supplementary Methods Paragraph 1.5.

### Molecular biology techniques

#### Western blotting

We performed western blotting using standard techniques. Samples were electrophoresed on SDS-PAGE gradient gels 4-20% (TGX stain free, BioRad) and proteins transferred onto nitrocellulose membranes (Bio-Rad). Membranes were blocked during 30 min with PBST-M (10 mM phosphate, 2.7 mM potassium chloride, 137 mM sodium chloride, pH 7.4 0.1% tween 20 (Sigma), 5% milk) and incubated with dilutions 1:1000 of anti-FLAG (Sigma, F1804 1*μ*g/*μ*L) overnight at 4°C (or 2h room temperature). Antibody binding was detected using the respective anti-mouseHRP secondary antibody (Proteintech) and Clarity ECL Western Blotting Substrate for chemiluminescence in a ChemiDocMP imaging system (BioRad).

#### UV crosslinking

For each strain, 100 ml bacterial cultures were grown to an OD_600_ of 2.0. Cultures were either directly irradiated in LB (no centrifugation step before irradiation) by placing on petri dishes which were kept on ice, exposure to UV light (λ = 254 nm) at 5 J/cm^2^ in a CL-1000 ultraviolet crosslinker device (Ultra-Violet Products Ltd) and centrifuged at 4°C for 10 min or centrifuged at room temperature for 10 min at 15,000 g and the pellets resuspended in 0.1 vol. of the original volume with water irradiated, and then centrifuged again (note that LB medium strongly absorbs UV light at 254 nm wavelength, resulting in inferior cross-linking efficiency).

#### Immunoprecipitation and PNK assay

Immunoprecipitation and PNK assay of bacterial FLAG-tagged proteins and radioactive labeling of RNA by PNK was performed as described (13). For details see Supplementary Methods Paragraph 1.5.

## Results

### TriPepSVM accurately recovers known RBPs with few false positives

We propose TriPepSVM, a SVM-based machine learning model to distinguish RNA-binding proteins from non-RNA binders based on the amino acid sequence of the protein of interest. To train our model, we collected the amino acid sequences of known RNA-binders (see Paragraph *Data sets*) as well as proteins that are very unlikely to bind RNA for both human and *Salmonella.* Figure 1 depicts how a protein-sequence is split into a set of overlapping k-mers. All of the k-mers of a sequence form a vector (see Paragraph *TriPepSVM prediction model*) which is then fed into the SVM classifier that learns a decision boundary to separate RBPs from non-RBPs.

**Fig. 1.**
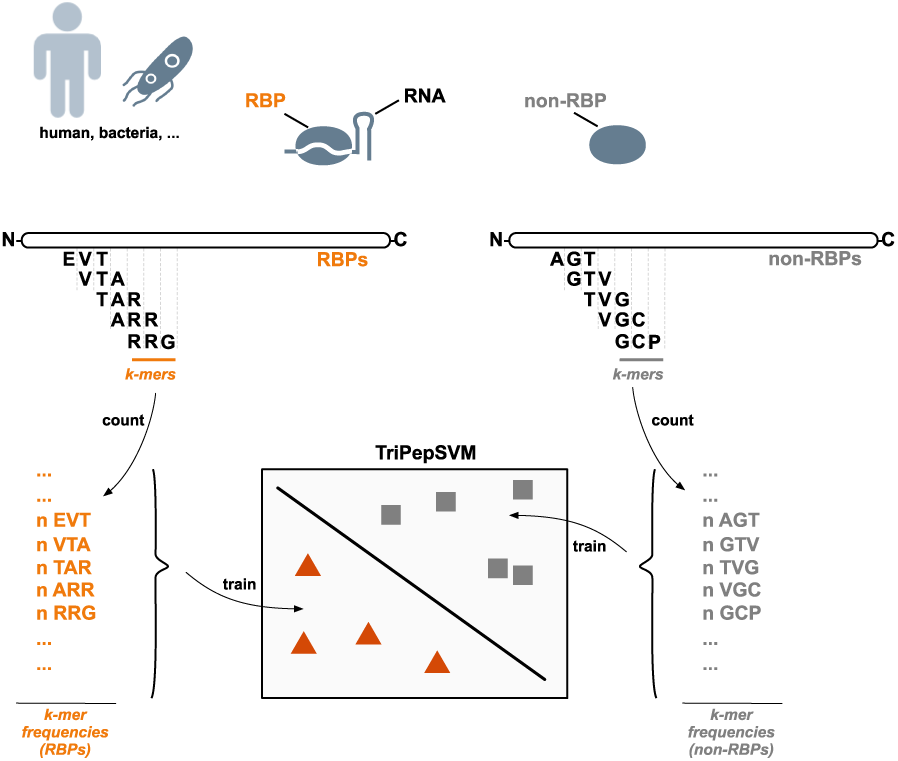
TriPepSVM schematic. TriPepSVM is a support vector machine trained on tri-peptide frequencies from RBPs and non-RBPs to discriminate between these two protien classes. It can be applied to different species.

Our proposed model has three hyper parameters that can heavily influence its performance. These are the parameter *C* that controls the penalty for mis-classifying data points, the k-mer length *k* of the spectrum kernel and finally the class weights of the positive and negative class (see Paragraph *Model training*).

**Table 1.**
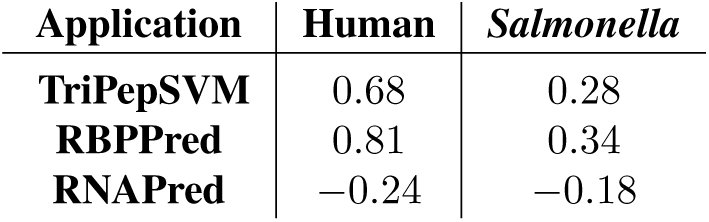
Optimal classification cutoff for the tested prediction tools.

After collection of the data, we therefore compute the best combination of hyper-parameters during model training by conducting a grid search on reasonable combinations of *C*, *k* and the class weights. Since we have only a very limited amount of known RBPs, we perform a 10-fold cross validation on the training data for each combination of hyper-parameters. The best combination is selected based on the best mean balanced accuracy. We select *C* = 1, *k* = 3 and class weights of 1.8 for the positive class and 0.2 for the negative class, respectively as best parameters for both, human and *Salmonella* (see Supplementary Figure S3 for more details on the parameter tuning results).

With the optimal combination of parameters at hand, we evaluated TriPepSVM on a held-out test set, which was neither used for training, nor hyper-parameter tuning (see Paragraph *Model training*). We show that TriPepSVM is capable of recovering most known RBPs in the test set while still maintaining a good specificity (not many false positives), yielding an area under the ROC curve (AUROC) of 0.83 and an Area under the Precision Recall Curve (AUPR) of 0.53 in human (see Figure 2). We compared the performance of TriPepSVM with formerly introduced methods for RBP prediction, namely SPOT-seq-RNA, RNApred and RBPPred and show that TriPepSVM outperforms all competing methods by a considerable margin (see Figure 2). Especially the PR curves show that TriPepSVM is far more precise than other methods, across all sensitivity cutoffs. SPOT-seq-RNA outputs a single classification result and no confidence score for it, hence its performance reduces to a single point on both of the curves.

**Fig. 2.**
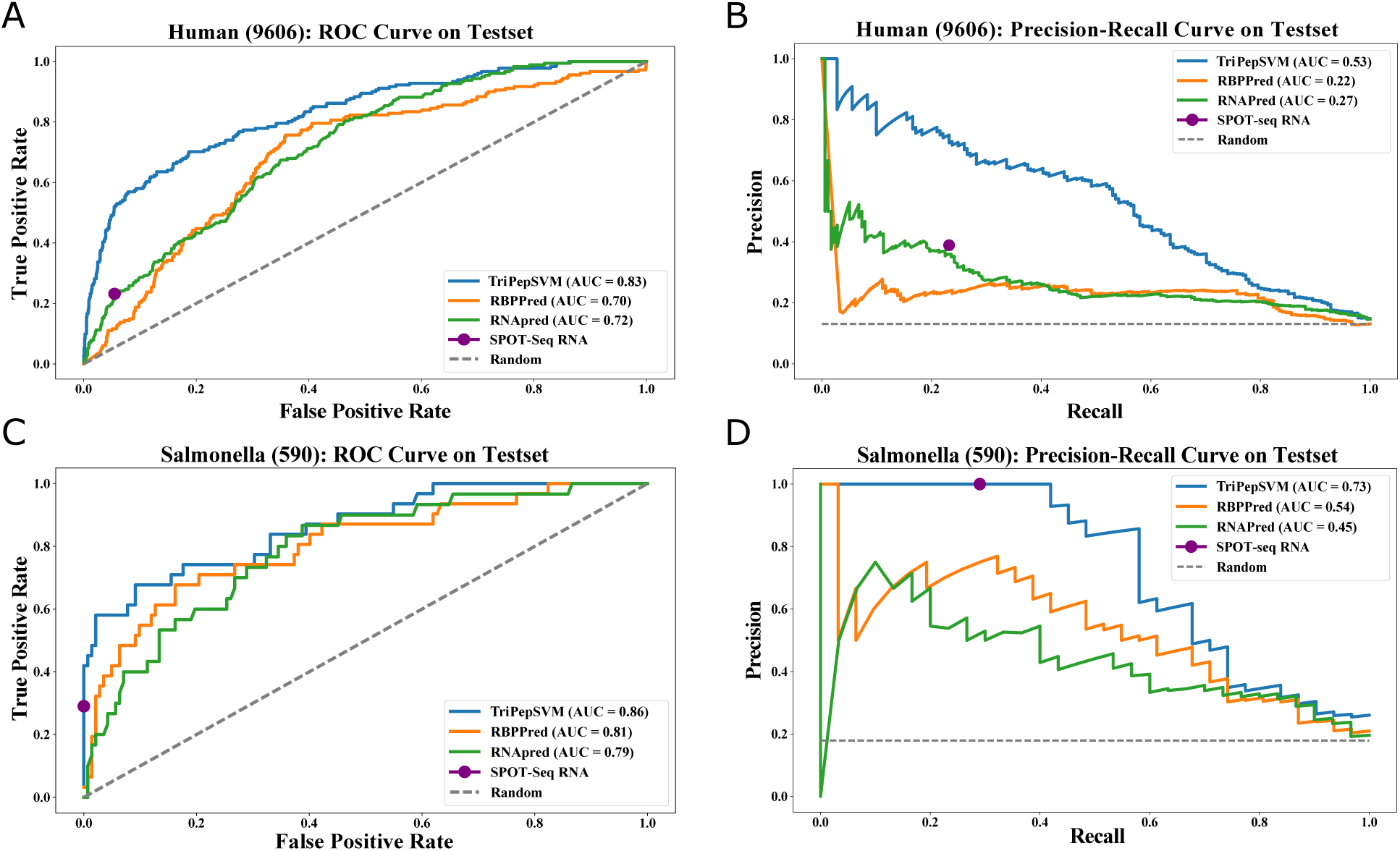
Performance of TriPepSVM in comparison to other RBP prediction methods. We compared our method TriPepSVM to RNAPred, RBPPred and SPOT-seq-RNA on both human and *Salmonella* test sets. **A & B** shows ROC and PR curves on the human test set. **C & D** shows ROC and PR curves on the *Salmonella* test set. SPOT-seq-RNA only outputs a class and no probability or score associated with the predictions and is hence represented only as a dot in the PR/ROC curves.

In addition, performance measures for all tools computed at the optimal PR curve cutoff (reported in Table 1 and determined as described in Supplementary Paragraph 1.4) are given in Supplementary Figure S4, for both human and *Salmonella.* TriPepSVM outperformed all other methods in terms of MCC on both human and *Salmonella* and had a slightly higher balanced accuracy compared to the best and most recent tool RBPPred. In addition, TriPepSVM reaches a good compromise between sensitivity and specificity at the optimal cutoff (Figure S4). In comparison, SPOT-Seq-RNA has a high specificity, as it is based on experimental RNA-RBP templates, but low sensitivity. On the other hand, RNAPred exhibits a sensitivity, relying only on amino acid composition, at the expenses of a poor specificity. The optimal cutoffs determined by TriPepSVM on both human and *Salmonella* were used to carry on proteome-wide predictions in both species.

### TriPepSVM results are consistent with interactome capture studies

We found 2944 proteins with a predicted RNA-binding capability in the human proteome (see Supplementary Table S11). To assess whether these proteins are really likely to bind RNA, we overlapped our predictions with interactome capture studies from recent years. Figure 3 shows the overlap between our predictions (TripPepSVM Predicted), the union of discovered RBPs by four different interactome capture studies and proteins that contain a Pfam RBD. The table in Figure 3 reports the fraction of proteins from the interactome capture studies which our model was able to recover for the tuned cutoff of 0.68. For all of the four studies, we were able to recover more than 75% of the experimentally identified RBPs, and for three of these studies this percentage was higher or equal to 85%.

**Fig. 3.**
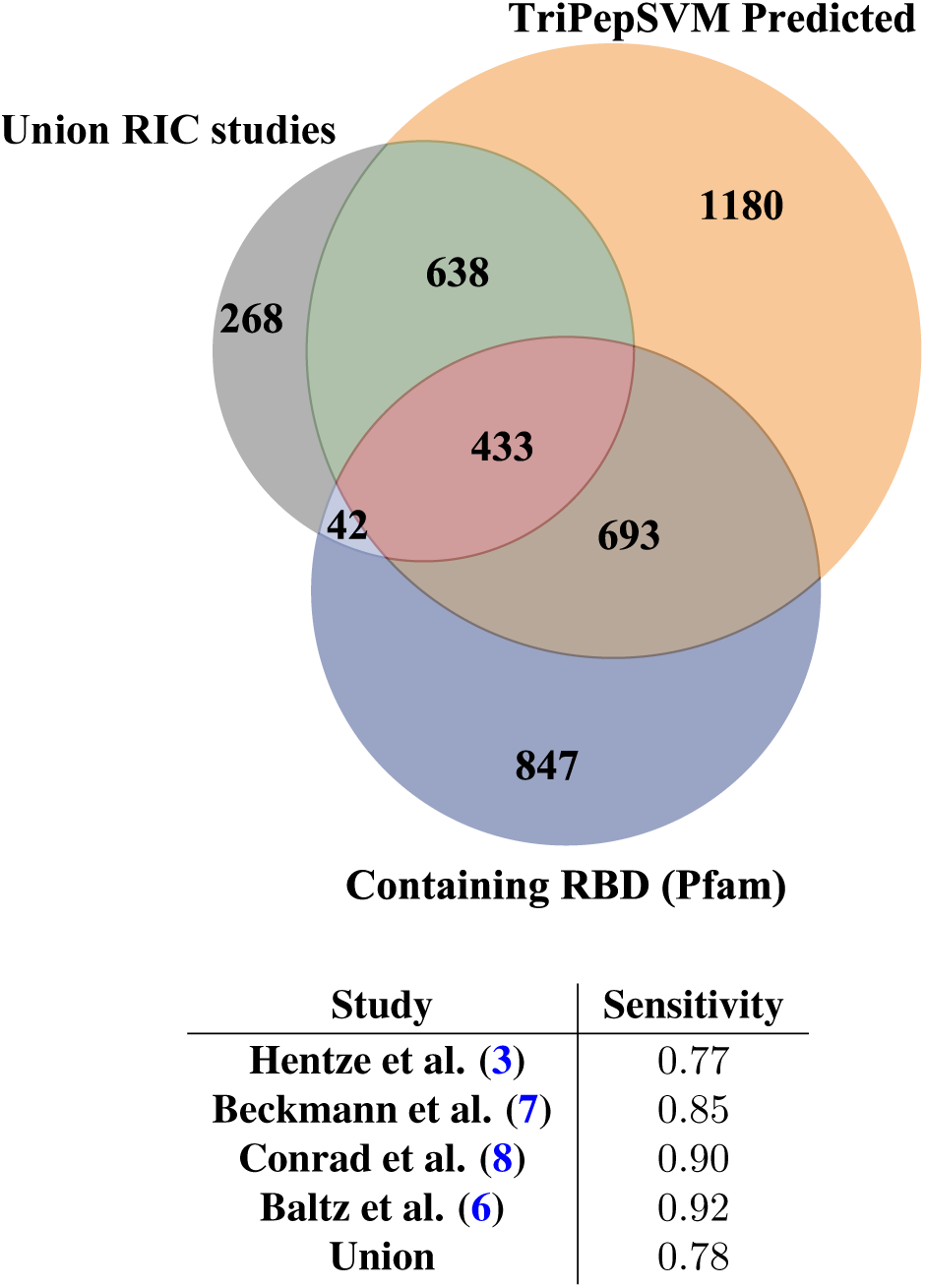
Overlap between four different interactome capture studies and predictions from TriPepSVM. We computed the overlap between our proteome-wide predictions (orange, top right), the union of identified proteins from four independent RIC studies (gray, top left) and proteins containing RBDs according to the Pfam database (blue, bottom). The table shows the sensitivity between our predictions and the four different RIC studies and their union.

### TriPepSVM predicts novel human RBPs

It should be taken into account that the vast majority of those RIC-RBPs are mRNA-binders since they were identified using poly-A RNA selection and eukaryotic mRNA only constitutes for a small subset of cellular RNA (around 5%). TriPepSVM however was trained on a superset of all known RBPs. Consistent with this, we also identify 14 cytosolic and 6 mitochondrial tRNA ligases, 6 tRNA/rRNA methyltransferases (NSUN2-6) and the majority of ribosomal proteins (31 out of 33 small subunit proteins and 44 out of 47 large subunit proteins (39)). The recent consensus for human RBPs ranges from hundreds (1) to more than 2000 (3) RBPs. From the 2944 proteins predicted by TriPepSVM, 990 (34%) were not previously described in the aforementioned RIC studies, do not contain a known RNA-binding domain or were not annotated as RBPs. Among the 990 predicted RNA-binders are more than 200 proteins with ATP-binding capacity, from these are 13 proteins of the AAA ATPase family. The overlap between RNA- and single nucleotide binding is generally very high among the established RBPs as well (8). Also enriched are kinases, proteins harboring WD40 domains and bromodomain folds. During the writing of this manuscript, we and others published preprints in which RBPs were identified by biochemical techniques (called PTex and XRNAX) in a poly-A independent fashion (40, 41). The findings in these studies are largely confirmatory to our results as proteins of the AAA ATPase family, WD40 and bromodomain proteins were likewise identified as novel RBPs.

### Important sequence patterns in RBPs and their biological significance

We next set out to identify those sequence k-mers or model features that contributed the most to classifying RBPs versus non-RBPs in both human and *Salmonella*, using Equation 2 (see Paragraph *Feature importance scores*). The highest ranked k-mers (top 50 ranking) are listed in the Supplementary Table S7 and Table S8 for human and *Salmonella*, respectively, while in Table S9 and Table S10 all k-mers with their corresponding weights are reported. In both organisms, k-mers containing lysine (K), arginine (R) and glycine (G) are found to have the largest SVM weight. Finding K and R enriched among the triplets was expected since positively charged residues are known to be involved in direct RNA interaction; e.g. the RGG box is a known RNA-binding motif in unstructured regions of proteins (11). Although an aromatic motif YGG has been identified as RNA-binding site by Castello and colleagues when mapping RNA-binding sites proteome-wide in human cells (11), in our dataset YGG is not among the top k-mers which contribute to RBP classification and it harbors an SVM weight value close to 0. YGG repeats (also called [G/S]Y[G/S] motif) can bind to RNA and promote hydrogel formation *in vitro* as well as liquid-liquid phase separations (LLPS) in vivo (42, 43). GYG, SYS, GYS and SYG triplets, which are potentially part of the [G/S]Y[G/S] motif, although not among the top 50 important k-mers in our human model, have all positive weights, indicating that they contribute to the recognition of the RBP class. Nearly all k-mers with a negative weight (contributing to classification as non-RBP) contain leucine (L) and/or glutamic acid (E). Consistent with this, E and L are the residues most absent from RNA-binding sites in human cells (11).

We then compared the weights of all k-mers from human and *Salmonella* to our previously identified triplets conserved in eukaryotic evolution (7) (see red triangles in Figure 4). Our findings independently confirm that those triplets are not only conserved among RBPs but are also important to correctly identify these proteins as RNA-binders by TriPepSVM, given that most of them have a positive weight (Figure 4). Interestingly, a high-portion of these k-mers found to be conserved in eukaryotic RBPs (mainly KR- and RG-containng triplets) seem to be important not only for human RBP, but also for bacterial RBP classification.

**Fig. 4.**
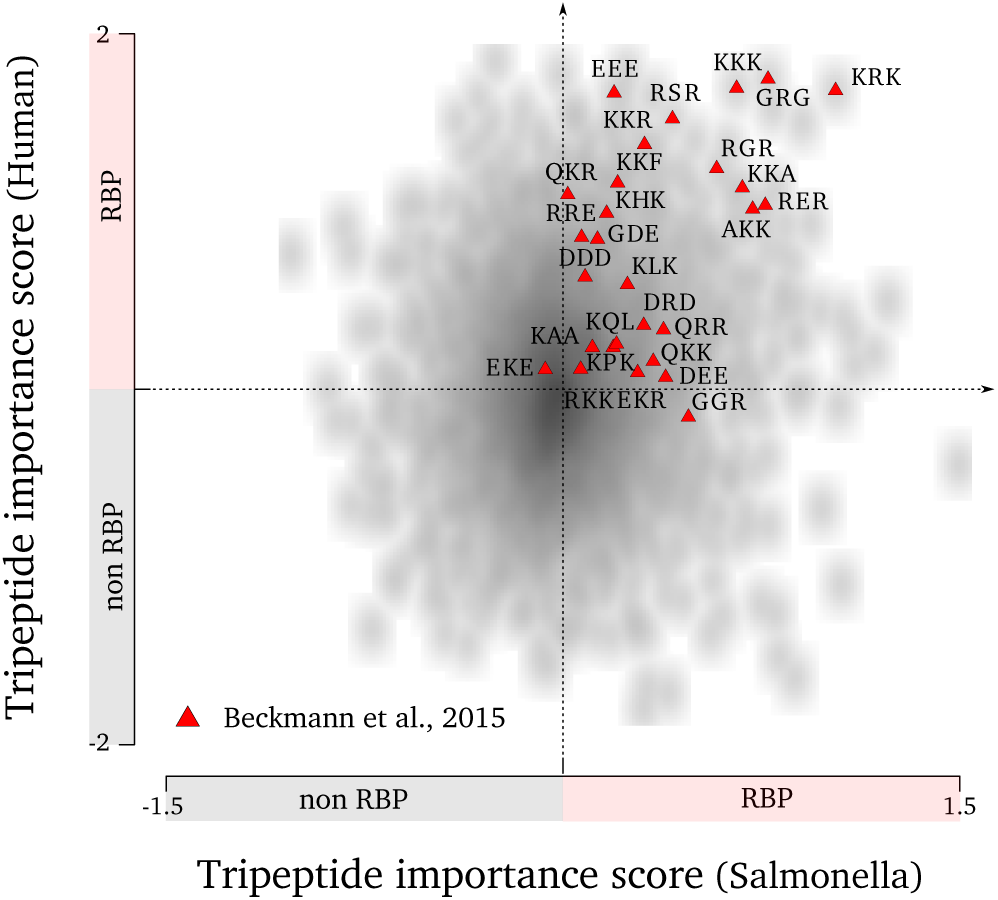
TriPepSVM classifiers apply highly conserved tripeptides for RBP classification. Shown are those tripeptides (red triangles) identified by Beckmann et al. (7) as conserved in eukaryotic RBPs and expanded from yeast to human with their corresponding weights from both human (y-axis) and *Salmonella* (x-axis) TriPepSVM classifiers. Most of the conserved tripeptides from (7) harbor positive weights not only in human but also in *Salmonella*, and are therefore important in characterizing RNA-binders in both species.

Following on the presence of k-mers which are known to bind RNA in unstructured regions, including those containing G/Y and R/G, we probed for both species whether overall the top k-mers identified by our SVM model had a higher propensity to be found in structured domains or unstructured parts of proteins (see Figure 5) using IUPred prediction (36). Strikingly, k-mers with the biggest contribution to classify a protein are more often found in disordered regions in human, but not in *Salmonella*. In addition, known human RBPs, such as HNRPU, PTBP1, FUS, SRSF1, U2AF2, DDX4 and others, implicated in several aspects of RNA processing, and known to mediate RNA binding via disordered regions, such as as R/G repeats, are correctly predicted by our method with very high probability (see Supplementary Table S11).

**Fig. 5.**
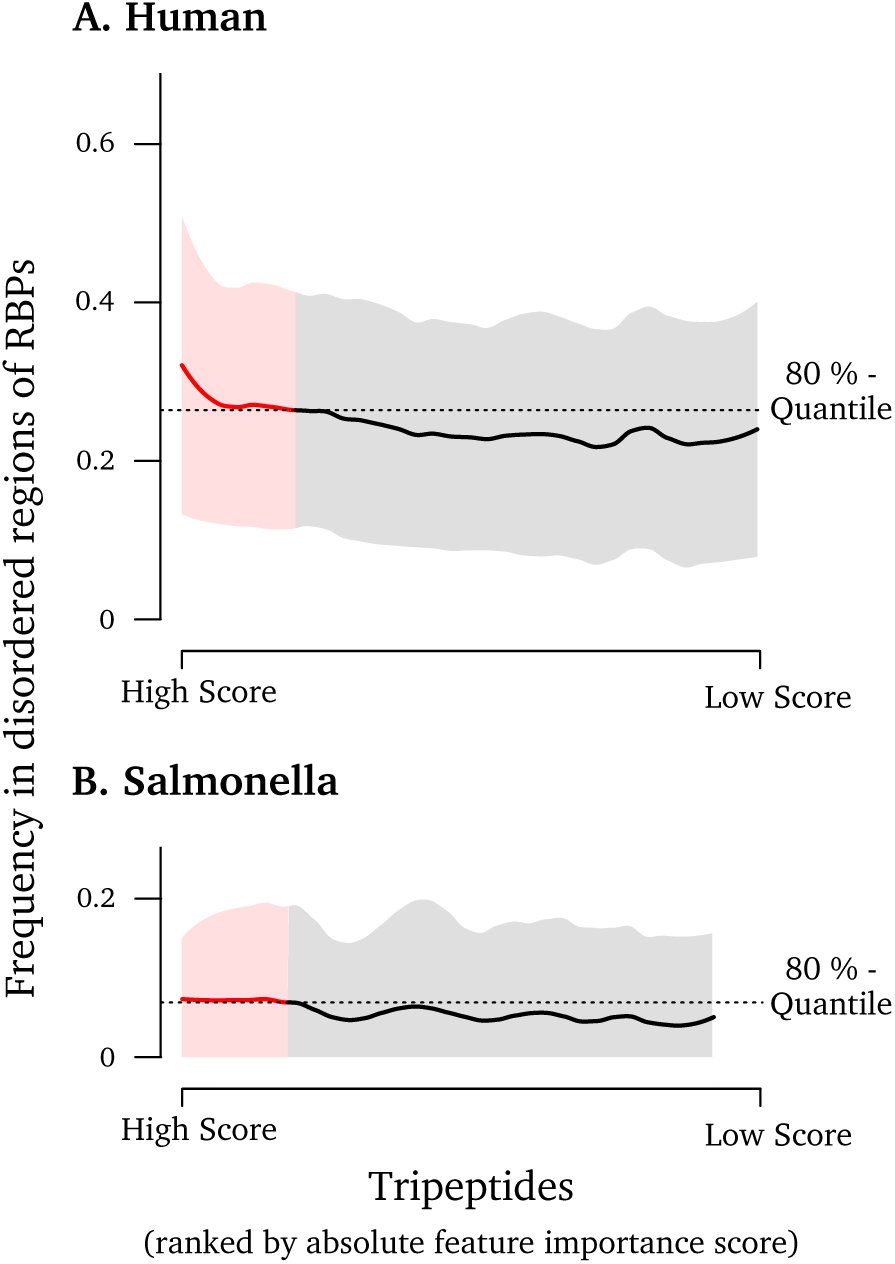
Important tripeptides are enriched in structurally disordered regions of the RBPs in human. The figure shows the structural disorder fraction of each tripeptide ranked by the absolute feature importance score. The resulting curve is smoothed using LOESS regression (span = 0.2) where the shading shows the standard deviation. The dashed line marks the value of the 80%-quantile of the smooth LOESS regression values. The frequency indicating how often a tripeptide is observed in the structural disordered part of an RBP increases with the feature importance score in **(A)** human, but not in **(B)** *Salmonella*.

### Searching the *Salmonella* proteome for RBPs

We next used TriPepSVM to predict potential RBPs in a prokaryotic organism. As for most bacteria, RBPs are poorly annotated in *Salmonella* Typhimurium despite recent advances such as the identification of the Csp proteins or ProQ as RNA-binders by Smirnov and colleagues. The same study however indicated that more, so far un-identified proteins harbor the potential to bind RNA (44). After training TriPepSVM on bacteria from the *Salmonella* clade (using the recursive mode for Unipro-tKB taxon 590) to obtain a larger positive training set, we searched the complete *Salmonella* Typhimurium proteome for RBPs. We correctly identify 108 of the Uniprot-annotated 115 RBPs, resulting in a 94% recovery rate. We correctly predict Hfq (13), and ProQ (44) which are bacterial mRNA- and sRNA-binding proteins. Only recently, cold shock proteins CspC/E were described to bind RNA (45). However, CspC is still not marked as RBP in the SwissProt Database and CspE was not present in the reviewed section of the database when training and testing TriPepSVM (see Paragraph *Methods*), explaining their absence from our predictions.

Using the tuned cutoff of 0.28 for *Salmonella* (see Table 1), we additionally predict 66 additional proteins to bind to RNA (see Supplementary Table S12). Among those are 8 ribosomal proteins and 14 other proteins involved in RNA biology all of which are not annotated as “RNA-binding” such as the GTPase Der which is involved in ribosome biogenesis (46), ribosomal methyltransferases RimO and RlmE (47), RNA pyrophosphohydrolase RppH (48), or transcription elongation factors GreA/B (49). Furthermore, we predict 20 known DNA-binding proteins and 18 proteins with documented ATP-binding activity to be RNA-interacting. Additionally, 18 predicted proteins are not implicated to interact with RNA or any other nucleic acid; of those 12 have enzymatic activity, consistent with a growing list of enzymes from diverse species to be associated with RNA (7, 10, 50).

### Experimental validation of predicted RBPs in *Salmonella*

Finally, we set out to experimentally validate TriPepSVM predictions *in vivo.* We generated *Salmonella* mutant strains carrying FLAG-tagged RBP fusion in its genomic context using the λ Red method; resulting in bacterial mutants that exclusively express the predicted FLAG-tagged RBP candidate at their respective physiological levels (38). We chose ClpX (a subunit of the Clp protease regulating expression of the flagellum (51)), DnaJ (a chaperone responding to hyperosmotic and heat shock (52)), UbiG (a ubiquinone biosynthesis O-methyltransferase (53)) and CysN (Sulfate adenylyltransferase subunit 1 (54)) as predicted RBPs and YigA which TriPepSVM predicts as non-RBP and we tested for RNA-binding *in vivo* using the PNK assay (see Figure 6A). As demonstrated by a radioactive signal from 5’ end labeled co-immunoprecipitated RNA (see Figure 6B,C), we can confirm that we correctly predicted ClpX, DnaJ, UbiG (RNA-binding) and YigA (true negative). The validation of 4 out of 5 proteins is therefore also matching with our calculated balanced accuracy of 73% (see Supplementary Figure S5). Importantly, ClpX has ATP-binding activity and DnaJ is able to bind to DNA and ATP (51, 52). To exclude self-phosphorylation or direct binding from ATP-binders to the radioactive isotope in our assay (55), we also included conditions in which PNK was omitted but P^32^-γ-ATP was provided (see Figure 6B,C). However, our validation demonstrates that both are bonafide RNA-interactors *in vivo* from which we conclude that TriPepSVM is unlikely to incorrectly predict DNA- or single nucleotide binders as RBPs.

**Fig. 6.**
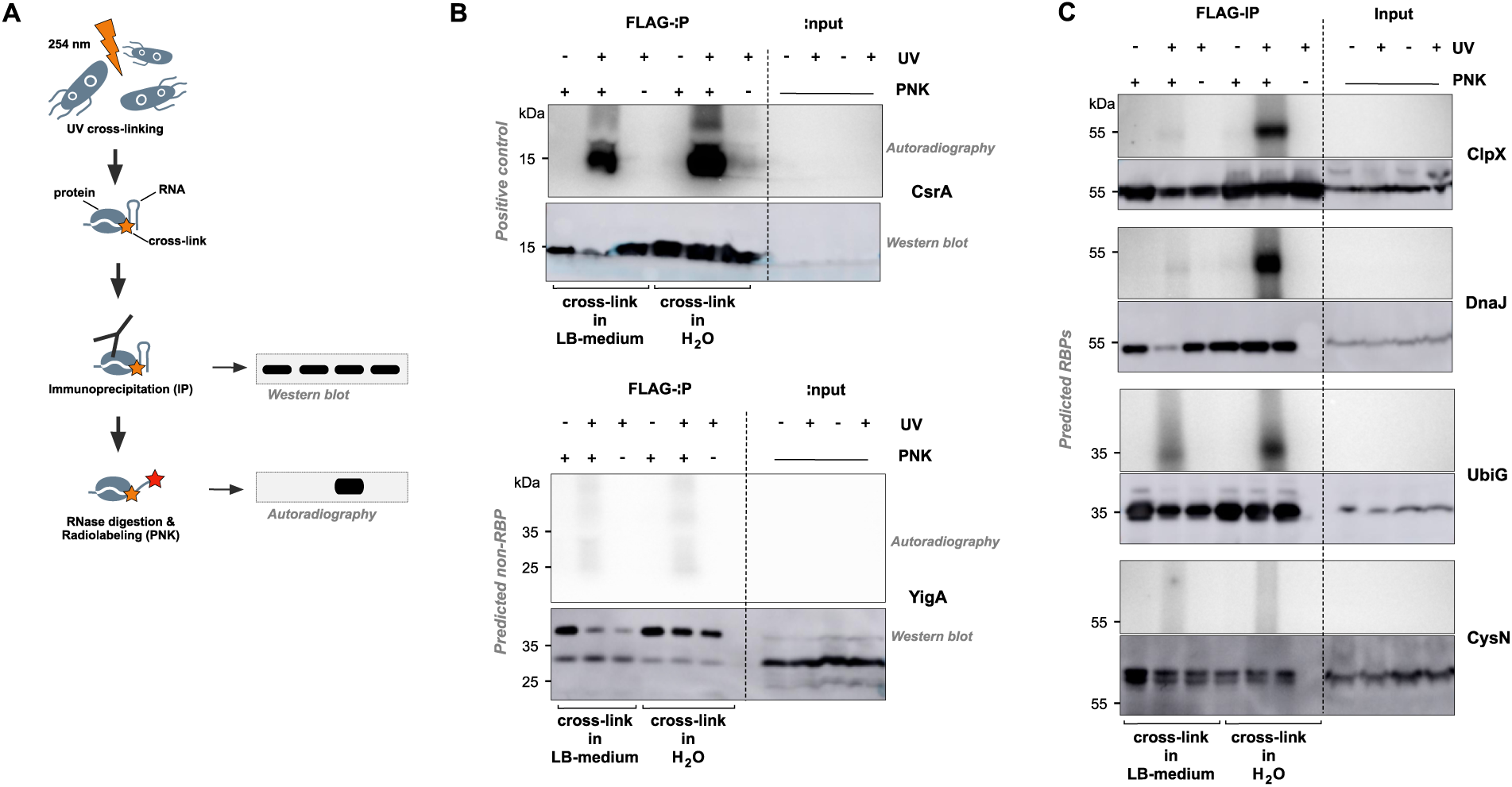
Experimental validation of predicted bacterial RBPs. **A** Schematic view of the PNK assay. After UV cross-linking of ribonucleoprotein complexes *in vivo*, cells were lysed. UV irradiation can result in a covalent link (orange star) between RNA and proteins that are in close physical proximity (zero distance cross-link). Individual candidate RBP-FLAG fusions were then immunoprecipitated (IP) with a FLAG-specific antibody. After trimming of the co-imunoprecipitated RNA using RNase digestion, polynucleotide kinase (PNK) was used to enzymatically add a radioactive phosphate (32P) to 5’-ends of transcripts. Finally, input controls as well as the IPs were separated by SDS-PAGE. After blotting to a membrane, protein amounts were analysed by Western blotting; presence of a radioactive signal at the same molecular mass as the protein then serves as indirect proof of RNA-binding. **B** PNK assay of CsrA (positive control) and YigA (predicted non-RBP). UV cross-linking was performed in LB medium and water independently. A radioactive signal can only be detected for RBPs (here: CsrA) after UV cross-linking and in presence of PNK. **C** PNK assay of 4 candidate RNA-binding proteins in *Salmonella* Typhimurium predicted by TriPepSVM. ClpX, DnaJ and UbiG could be confirmed to interact with RNA *in vivo*, but not CysN.

## Discussion

What defines a RNA-binding protein? Apart from the obvious functionality to bind to RNA, other elements within a protein can be important to exert its physiological role in the cell. With TriPepSVM, we are presenting an approach which reduces a protein to its combination of short amino acid k-mers, in our case triplets, and use machine learning to find patterns in these combinations that align with RNA-binding. *Salmonella* Typhimurium is a well-studied bacterium; not the least due to its role as Gram-negative model organism to study infection by prokaryotic pathogens. Despite its importance, only a very limited set of RNA-interacting proteins has been identified in *Salmonella* and other bacteria beyond the canonical set of proteins which make up the transcription machinery, the ribosome or interact with tRNA. In recent years, novel approaches such as Grad-Seq (44) identified additional proteins that can interact with bacterial mRNA. So far, 5 mRNA-binding proteins have been confirmed in *Salmonella*: Hfq, CsrA, ProQ and CspC/E. This limited data set renders prediction and discovery of novel RBPs in bacteria a challenging task since most bioinformatic prediction tools depend on either structural similarity to protein folds that are known to be involved in RNA interaction or on homology based on phylogeny. In both cases, the limited available data set of known bacterial RBPs represents an important obstacle - also for our method. Still, our approach correctly identifies most known RNA-binders and predicts 66 novel candidate RBPs in *Salmonella* from which we tested ClpX, DnaJ, UbiG and CysN for validation. Indeed, 3 (ClpX, DnaJ and UbiG) out of the 4 could be confirmed to bind RNA *in vivo* (see Figure 6). In our approach, we reduce the search space to the most basic feature of any protein: its primary sequence. Following the observation that i) many recently-found RBPs lack known RNA-binding domains (11) and ii) our earlier work showed an expansion of short triplet amino acid motifs in RBPs throughout evolution (7), TriPepSVM rather searches for combinations of triplet peptides in proteins then for full domains. This reduction in complexity has the advantage that TriPepSVM is independent on prior (and potentially biased) knowledge on RBDs or homology.

The fact that TriPepSVM does not classify all human proteins with a Pfam-domain (see Figure 3) as RBP also demonstrates that tripeptides from RNA-binding domains alone are not sufficient to explain the performance of TriPepSVM. Intriguingly, tripeptides which we predict to contribute to RNA-interaction more prominently have a tendency to be enriched in structurally disordered regions in human but not in *Salmonella* (see Figure 5). Together with our earlier comparison of tripeptide motifs in eukaryotic RBPs in which unicellular yeast harbors few tripeptide repetitions that expand during evolution, it is tempting to speculate that RNA-binding via unstructured regions is of higher physiological relevance in more complex organisms than in unicellular species. Consistent with this hypothesis is the observation that sequence-independent RNA-binding in unstructured regions of RBPs is important for P-bodies or RNA granules by liquid-liquid phase transitions (56). Formation of these higher-order RNA-protein complexes however has not been described for bacteria so far. Our results show that if present in prokaryotes, regulation of RNA-granule-like complexes is very unlikely through unstructured regions of RBPs.

## Conclusion

All in all, we show that the propensity of a protein to bind RNA is mostly encoded in its primary sequence and can be confidently predicted based solely on combinations of short amino acid triplets. TriPepSVM outperforms previous approaches which make use of more complex protein features in discriminating RBPs from non-RBPs. It can in principle be applied to any species, from eukaryotes to bacteria where limited experimental data are available. Besides being a valuable RBP prediction method from sequence alone, our approach can pinpoint the important sequence patterns which distinguish RBPs from non-RBPs and points to disordered regions as main determinants of RBP-RNA interactions, in line with the latest studies.

## Data Availability

The collection pipeline as well as the source code for TriPepSVM are available on Github, https://github.com/marsicoLab/TriPepSVM.

## Acknowledgements

Work in the Marsico lab is supported by the DFG Grant MA 4454/3-1. Work in the Beckmann lab is supported by the DFG (IRTG 2290 and ZUK 75/1 Project 0190-854599). AB and RSS are supported by the International Max Planck Research School for Computational Biology and Scientific Computing. ECU is supported by the Joachim Herz Foundation. We are thankful to Jörg Vogel for providing plasmid pSUB11 and *Salmonella* strains, to Barbara Plaschke and Jens Hör for help with generating the mutants, and to Julie Bohl for technical assistance.

